# Telomere length, arsenic exposure and risk of basal cell carcinoma of skin

**DOI:** 10.1101/465732

**Authors:** Nalini Srinivas, Sivaramakrishna Rachakonda, Thomas Hielscher, Silvia Calderazzo, Peter Rudnai, Eugen Gurzau, Kvetoslava Koppova, Tony Fletcher, Rajiv Kumar

## Abstract

Telomere length *per se* a heritable trait has been reported to be associated with different diseases including cancers. In this study based on arsenic exposed 528 cases with basal cell carcinoma of skin (BCC) and 533 healthy controls, we observed a statistically significant association between decreased telomere length and increased BCC risk (OR = 5.92, 95% CI = 3.92-9.01, *P*<0.0001). We also observed that higher arsenic exposure (>1.32 µg/L) was statistically significantly associated with decreased telomere length (β = −0.026, 95% CI = − 0.05-0.003, *P* = 0.02). The interaction between arsenic exposure and telomere length on BCC risk was statistically significant (*P* = 0.02). Within each tertile based on arsenic exposure, the individuals with shorter telomeres were at an increased risk of BCC, with highest risk being in the highest exposed group (OR = 16.13, 95% CI = 6.71-40.00, *P*<0.0001); followed by those in medium exposure group (OR = 5.05, 95% CI = 2.29-10.20, *P* <0.0001), and low exposure group (OR = 3.44, 95% CI = 1.81-6.54, *P* = 0.0002). The combined effect of highest arsenic exposure and shortest telomeres on the risk of BCC (OR = 10.56, 95% CI = 5.14-21.70) showed a statistically significant departure from additivity (interaction constant ratio 6.56, *P* = 0.03). Our results show that in the presence of arsenic exposure, decreased telomere length predisposes individuals to increased risk of BCC, with the effect being synergistic in individuals with highest arsenic exposure and shortest telomeres.

## Introduction

Basal cell carcinoma (BCC) of skin, which arises from the transformation of keratinocytes within the epidermis, accounts for 80-90% of all primary skin cancers. BCC are slow-growing and locally invasive tumors that result in extensive morbidity through recurrence and tissue destruction (1). In genetically predisposed populations, ultraviolet radiation through sun exposure constitutes a major risk factor for BCC. Other risk factors include ionizing radiation, chronic inflammatory skin conditions, and arsenic exposure (2,3). BCC occurs mainly sporadically but some rare genetic disorders like Gorlin’s syndrome and xeroderma pigmentosum result in multiple tumors with early age of onset (4).

Telomeres at the chromosomal ends in humans consist of TTAGGG repeat sequences of approximately 10-15 kb of double stranded DNA ending in a single strand of up to 150-200 nucleotides (5). Telomeres together with shelterin complex proteins afford protection from end fusions to maintain genomic integrity (6,7). Telomere repeats, in the absence of telomerase in most of the somatic cells, progressively shorten by ∼200 nucleotides with each cell division due to the ‘end-replication problem’ (8,9). Short or dysfunctional telomeres are recognized as DNA double stand breaks, triggering cells to undergo replicative senescence. Telomere length is considered as a potential biomarker of aging and age-related diseases (5). Studies conducted in large cohorts have reported that longer telomeres are associated with increased risk of different cancers, including melanoma, BCC, lung cancer, gliomas, lymphoma, and bladder neoplasms (10,11).

Telomere length is also influenced by a wide spectrum of genetic and environmental factors, which include various inherited syndrome, collectively called telomeropathies, electromagnetic radiation, physiological stress, lifestyle factors and exposure to carcinogens such as arsenic (12-15). Arsenic exposure induces oxidative stress leading to DNA damage, genomic instability, and shortened telomeres (16-18). Epidemiological studies have indicated that the long-term arsenic exposure causes increased risk of cancers of the skin, bladder, lung and kidney (19,20). However, only a few studies have focused on the effects of arsenic exposure on telomere length with limited subjects (21,22). In the present multi-center study, we observed that the BCC patients with different life time arsenic exposure had statistically significantly shorter telomeres than the healthy controls.

## Methods

### Study Population

This study constituted a part of a multinational European research project conducted in several regions of Hungary, Romania and Slovakia between 2002 and 2004. The cases and controls selected were of Hungarian, Romanian and Slovakian nationalities. On the basis of histopathological examinations by pathologists, 528 BCC cases and 533 hospital-based controls were included in the study, subject to fulfillment of a set of criteria as described previously (23). The blood samples were kept at −80°C until analysis. A general questionnaire that included cumulative sun exposure, effects of sun exposure on skin and age/s at diagnosis of skin BCC was completed by trained personnel after an interview of the recruited cases and controls. In addition, the interviews included details on demography, life-style, socio-economic, medical history, occupational exposures, drinking and nutritional habits as well as detailed residential history (3,19,24). The concentration of arsenic in drinking water over the lifetime of each individual had been measured in the study population. The assessment of arsenic exposure through drinking water has been described previously (25). Ethnic background of the cases and controls were recorded along with other characteristics of the study population. Local ethics boards approved the study plan and design.

### Measurement of Relative Leukocyte Telomere Length

Relative leukocyte telomere length was measured in DNA from 528 BCC cases and 533 healthy controls using a multiplex quantitative real time PCR based assay, as described earlier by Cawthon et al., with minor modifications (26,27). Briefly, each reaction was performed in triplicates in optical 384-well reaction plates, in a 10µl volume, using 2µl of 5X HOT FIREPol probe qPCR Mix Plus with ROX (Solis BioDyne, Tartu, Estonia), 1.5µM of Syto-9 (Invitrogen, Thermo Fischer Scientific Inc., USA) and 5-10ng of genomic DNA. Non-template controls without any genomic DNA were included as negative controls. Four primers were used in each reaction to amplify telomere DNA (telg at 200nM and telc at 400nM) and the albumin gene (albugcr2 at 200nM and abldgcr2 at 400nM) and the primer sequences were designed accordingly **(Supplementary Table S1)**. Real-time PCR experiments were performed on a Viia-7 instrument (ABI, Applied Biosystems, Foster City, CA) using two simultaneous programs to acquire the respective C_T_ values for telomere sequence and albumin gene. The conditions for amplification of the telomere sequence were 95°C/15 min, 2 cycles of 95°C/20s and 49°C/1 min followed by 27 cycles of 85°C/20s with signal acquisition at 59°C/30s. The conditions for albumin gene were 35 cycles of 95°C/15s, 85°C/30s with signal acquisition at 84°C/30s. The specificity for all amplifications was determined by melt-curve analysis performed at default settings (95°C/15s, 60°C/1min with continuous signal acquisition at 0.05°C/s ramping and 95°C/15s). Eight concentrations of a reference DNA sample (genomic DNA pooled from 17 healthy individuals) were included in triplicates in a 2-fold serial dilution (from 50ng to 0.4ng) to generate standard curves for telomere (T) and albumin (S) PCR products. The quality of the PCR amplification was assessed using Applied Biosystems Viia-7 RUO software, version 1.2.2. The standard curve was used to quantify the telomere and albumin genes based on their respective C_T_ values and the obtained triplicate values were averaged. The relative telomere lengths were expressed as the ratio between T and S (T/S ratio). Inter-assay variation and intra-assay variation was determined by duplicating the reference DNA at all the dilutions in each of the assay. The inter-assay coefficient of variation (CV) of the telomere and albumin assays were 1.52% and 0.90%, respectively and the intra-assay CV of the telomere and albumin assays were 0.63% and 0.67%, respectively. The PCR efficiencies of the telomere and albumin gene assays ranged between 98% and 103%.

### Statistical analysis

The telomere length as a continuous variable was log-transformed. Box plots were drawn to show the distribution of telomere length (represented by T/S ratio) in BCC cases and controls and the differences were analyzed by two-tailed t-test. Scatter plots were drawn using telomere length as dependent variable and age as independent variable. Regression lines were drawn to summarize the correlation between the variables. The regression model that included the covariates was given by the equation: telomere length = ß0+ß1*age +ß2*BCC status +ß3*age*BCC status for each subject. The confidence intervals for beta coefficient/slope and mean differences between telomere length in cases and controls, and corresponding *P* values were estimated using generalized linear regression model by adjusting with sex as a co-variate.

As a categorical variable, the telomere length was grouped into three based on tertile distribution of T/S ratio (q1 = ≤ 0.49, q2 = 0.50-0.73, q3 = >0.73). The association of telomere length and other co-variates that included age, sex, eye color, country, skin complexion, skin response to sun exposure, arsenic exposure, rs861539 (*XRCC3*) and *MC1R* polymorphisms with BCC risk was estimated using univariate unconditional logistic regression. The risk estimates were also adjusted using a multivariate logistic regression and the corresponding odds ratios (OR) and 95% confidence intervals (CI) were determined.

Mendelian randomization (MR) was implemented to determine the causal effect of telomere length (exposure) on the risk of BCC (outcome) using genetic variants as instrumental variables. The Mendelian randomization was based on the following assumptions: i) the genetic variants are associated with the exposure ii) the variants are independent of any confounders of the exposure-outcome association iii) the variants are not associated with the outcome conditional on the exposure and confounders of the exposure-outcome relationship (28). Mendelian randomization was carried out in R v3.4.3 (R project for statistical computing) using the R-package ‘Mendelian Randomization’ (29). In this model, the risk estimates were determined using inverse-variance weighted (IVW) and maximum likelihood-based methods. Nine single nucleotide polymorphisms (SNPs) as valid instrumental variables were chosen from previously published genome-wide association studies (30-32). In a two-sample MR study, the ß-estimates and the standard errors of the instrument-exposure association were taken from the genome-wide association studies (30-32). The associations between the instrument-outcome were estimated using logistic regression at the allele levels in the study population. A one-sample MR study was conducted using non-overlapping datasets in the same population (33). Two datasets were generated for cases and controls based on similar mean and variance for arsenic exposure. The associations of the instrument-exposure were estimated using linear regression analysis in the combined population and the control population of the first N/2 individuals (dataset 1). Linear regression model included telomere length as dependent variable with polymorphism, age (continuous), arsenic exposure (continuous), country and polymorphism*age as independent variables. The standardized beta estimates and standard error values for the variant allele of each polymorphism*age on telomere length was estimated. The associations of the instrument-outcome were estimated using logistic regression, adjusting for the covariates-age, arsenic exposure and country, in the second N/2 individuals (dataset 2).

Linear regression model was carried out to determine the effect modification of log-transformed arsenic exposure on log-transformed telomere length. The analysis was carried out using telomere length as a dependent variable and arsenic exposure, age and country as independent variables in the combined population. Arsenic exposure was taken as a continuous variable and in individual categories based on the median distribution, with ≤1.32 µg/L as lower exposure group and >1.32 µg/L as higher exposure group. Arsenic exposure was also modeled as a continuous variable based on WHO’s guideline for arsenic in drinking water (≤10 µg/L and >10 µg/L).

Further, the interaction between telomere length (log-transformed continuous variable) and arsenic exposure (log-transformed continuous variable) on BCC risk (as dependent variable) was assessed. Arsenic exposure was categorized based on tertile distribution (q1 = ≤0.70, q2 = 0.71-16.38, q3 = >16.38). Univariate logistic regression analysis for telomere length (as a continuous variable) on the risk of BCC was conducted in each of the three tertiles of arsenic exposure. Telomere length was adjusted for all the significant confounders in a multivariate regression analysis on the risk of BCC in each tertile of arsenic exposure.

Arsenic exposure-telomere length interaction on BCC risk was estimated by testing departure from additive or multiplicative odds-ratios. Based on tertile distribution, arsenic exposure was categorized as high exposure (HE), medium exposure (ME) and low exposure (LE) groups; and telomere length was categorized as long (L), medium (M) and short (S). The odds ratios were determined using logistic regression analysis adjusting for country and individuals age. Lowest exposure and longest telomere length, R_HE_R_L_ was used as reference to calculate the odds ratios for the remaining 8 groups. The estimates were calculated as R_HE_R_S_ for high exposure and short telomere length, R_HE_R_M_ for high exposure and medium telomere length, R_HE_R_L_ for high exposure and long telomere length; R_ME_R_S_ for medium exposure and short telomere length, R_ME_R_M_ for medium exposure and medium telomere length, R_ME_R_L_ for medium exposure and long telomere length; R_LE_R_S_ for low exposure and short telomere length and R_LE_R_M_ for low exposure and medium telomere length. Interaction constant ratio (ICR) was calculated to test departure from additivity (< or > 0) for high exposure group and short telomere length as (ICR_high_= R_HE_R_S_ - R_HE_R_L_- R_LE_R_S_+1) and high exposure group and medium telomere length as (ICR_high_= R_HE_R_M_ - R_HE_R_L_- R_LE_R_M_+1). Multiplicative interaction index (MII) was calculated to test departure from multiplicativity (< or > 1) for high exposure group and short telomere length as (MII_high_= R_HE_R_S_ / (R_LE_R_S_*R_HE_R_L_) and high exposure group and medium telomere length as (MII_high_= R_HE_R_M_ / (R_LE_R_M_*R_HE_R_L_). Similarly, ICR for medium exposure group and short telomere length was calculated as (ICR_med_= R_ME_R_S_ – R_ME_R_L_- R_LE_R_S_+1) and medium exposure group and medium telomere length as (ICR_med_= R_ME_R_M_ – R_ME_R_L_- R_LE_R_M_+1). MII for medium exposure group and shortest telomere length as (MII_med_= R_ME_R_S_ / (R_LE_R_S_*R_ME_R_L_) and medium exposure group and medium telomere length as (MII_med_= R_ME_R_M_ / (R_LE_R_M_*R_ME_R_L_). Confidence intervals and *P*-values for ICR and MII were determined using bootstrap method with 10,000 simulations. Statistical analysis with the exception of Mendelian randomization was performed using SAS, v.9.4 (SAS Institute, Inc., Cary, NC).

## Results

### Telomere length and BCC risk

The median age of 528 BCC cases at diagnosis was 67 years (Interquartile range, IQR = 58-73 years) and that of 533 controls was 61 years (IQR = 52-70 years) with 236 (44.70 %) men and 292 (55.3 %) women in cases and 274 (51.4 %) men and 259 (48.6 %) women in controls. Relative telomere length was successfully measured in 524 cases and 527 controls by multiplex quantitative real-time PCR. Analysis of the data showed that the BCC patients carried statistically significantly shorter telomeres than the controls (T-test, *P* = 6.3 x 10^-19^). The median log transformed telomere length in the cases was -0.58 (IQR = −0.82 to -0.40) and in the controls -0.39 (IQR = −0.58 to -0.25). A statistically significant inverse correlation (Pearson’s correlation r -0.30, *P*<0.0001) was observed between telomere length and age. The telomere attrition per year was similar both in the cases (slope = −0.007, 95% CI = −0.01 to -0.005, *P*<0.0001) and the controls (slope = −0.007, 95% CI = −0.01 to -0.005, *P*<0.0001) and the mean difference in telomere attrition per year between the two groups, adjusted for sex, was not statistically significant (estimate = −0.0003 95% CI = −0.004 to 0.003, *P* = 0.86; **Figure 1**).

**Figure 1:**
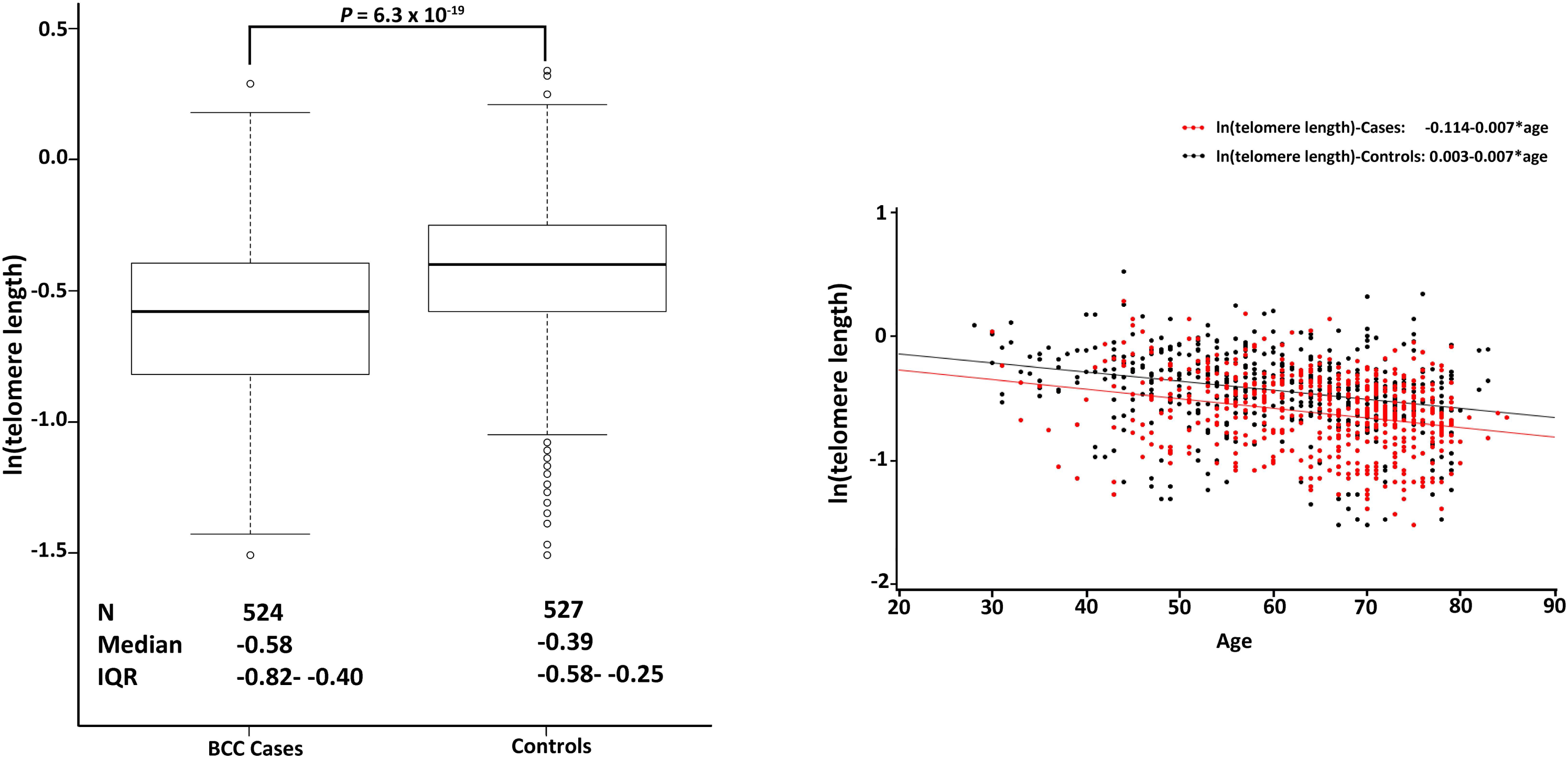
Distribution of ln(telomere length) in cases and controls. The box plots on the left depict differences in ln(telomere length) between BCC cases and controls. Number of cases and controls (N), median telomere length (median) and the corresponding interquartile range (IQR) are underneath the plots. Scatter plots on the right show relationship between ln(telomere length) and age in cases (red dots) and controls (black dots). Regression lines and corresponding equations are shown on the top right.

**Figure 2:**
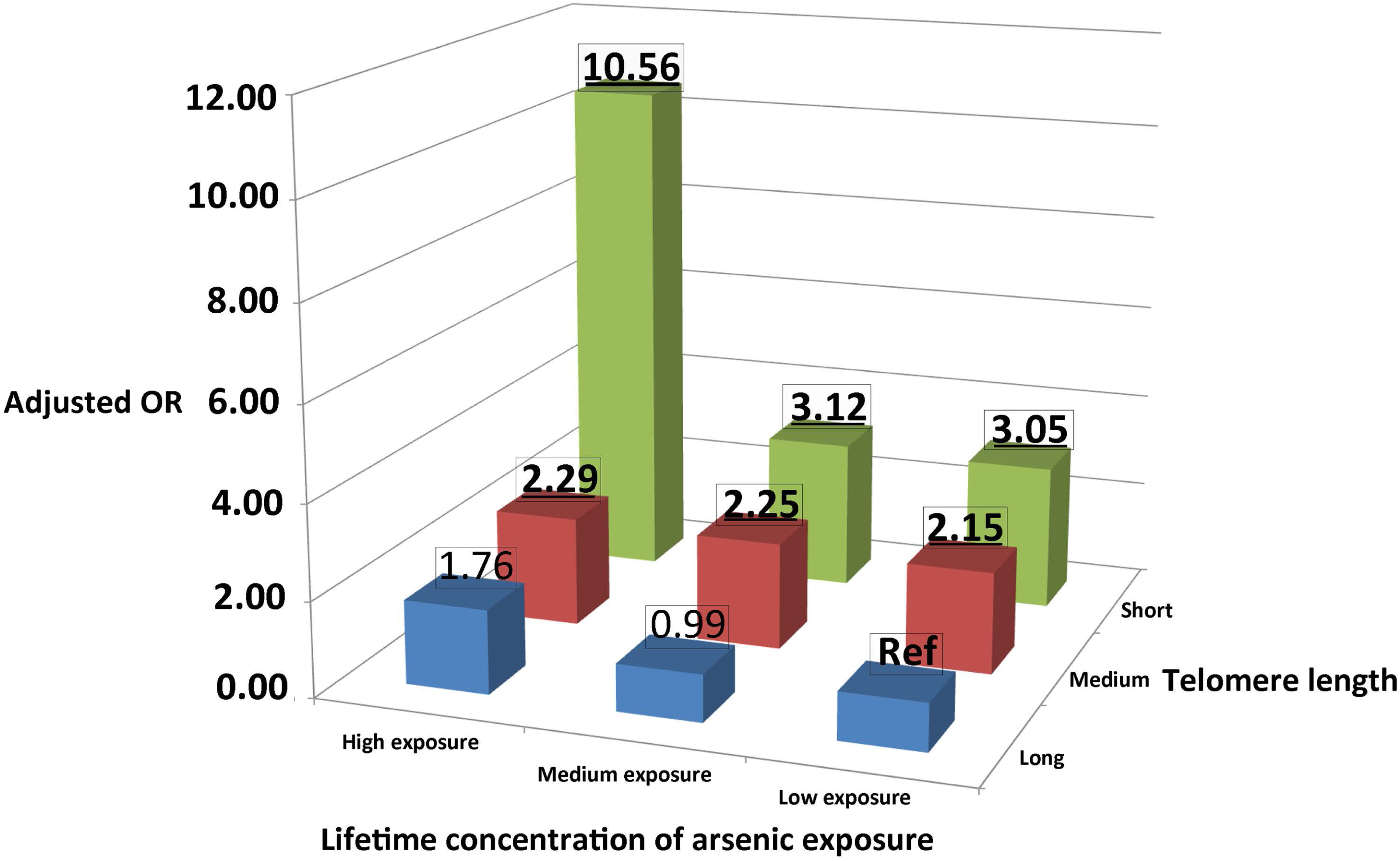
Risk of BCC associated with the combined effect of arsenic exposure and telomere length. OR associated with BCC risk due to various combinations is given on top of bars and statistically significant values, indicated in bold and underlined. Lifetime concentration of arsenic exposure was categorized as high, medium and low exposure. Telomere length was categorized as short, medium and long. The combination with lowest arsenic exposure and longest telomeres was used as reference. The interaction between the highest arsenic exposure and shortest telomere length was greater than additive.

A logistic regression model was used to determine the association between telomere length, both as a continuous and categorical variable, and risk for BCC. The effect of each unit decrease in telomere length on BCC risk was estimated to have an OR of 5.92 (95% CI = 3.92- 9.01, *P*<0.0001). Other factors that associated with BCC risk included age, sex, eye color, skin complexion, skin response to sun exposure, country and life-time arsenic exposure (taken as log transformed). We also included the data on melanocortin 1 receptor (*MC1R*) variants and the rs861539 polymorphism in *XRCC3* in the analysis that were previously shown to be associated with BCC risk in the current study population (23,34) **(Supplementary material and Supplementary Table S2)**. The OR for the effect of decreased telomere length on the risk of BCC after adjustment with all confounders was 4.57 (95% CI = 2.82-7.41, *P*<0.0001; **Table 1)**. Analysis of telomere length as categorical variable based on tertile distribution showed that individuals in the first tertile (shortest telomeres) compared to those in the third tertile (longest telomeres) carried largest risk for BCC (OR = 4.74, 95%CI = 3.46-6.50), followed by individuals in second tertile (OR = 2.06, 95%CI = 1.52-2.80), *P*<0.0001 **(Supplementary Table S2)**.

**Table 1.** Multivariate Analysis for association between telomere length and BCC risk

|  |  |  |  |  |  |
| --- | --- | --- | --- | --- | --- |
| **Effect** |  | **Cases** | **Controls** | **OR (95% CI)** | **P-value<sup>b</sup>** |
| **Telomere length<sup>a</sup>** | *Continuous (log-transformed)* | 480 | 490 | **4.57 (2.82-7.41)** | **<0.0001** |
| **Age** | *Continuous* | 480 | 490 | **1.03 (1.02-1.04)** | **<0.0001** |
| **Sex** | Male | 217 | 251 | **Ref** | 0.06 |
|  | Female | 263 | 239 | 1.31 (0.99-1.74) |  |
| **Eye Color** | Blue/Green | 290 | 260 | 1.26 (0.93-1.70) | 0.18 |
|  | Hazel/Brown | 164 | 201 | **Ref** |  |
|  | Others | 26 | 29 | 0.81 (0.43-1.53) |  |
| **Country** | Hungary | 145 | 230 | **Ref** | **<0.0001** |
|  | Romania | 129 | 129 | **2.50 (1.53-4.09)** |  |
|  | Slovakia | 206 | 131 | **3.48 (2.20-5.51)** |  |
| **Arsenic exposure (in µg/L) (log-transformed)** | q1: ≤0.70 (≤-0.36) | 166 | 150 | **Ref** | 0.06 |
|  | q2: 0.71-16.38 (-0.35- 2.80) | 166 | 165 | 1.16 (0.81-1.67) |  |
|  | q3: >16.38 (>2.80) | 148 | 175 | **1.82 (1.11-2.99)** |  |
| **Skin complexion** | Light | 257 | 198 | **1.69 (1.25-2.28)** | **0.0007** |
|  | Medium/Dark | 223 | 292 | **Ref** |  |
| **Skin response to sun exposure** | Blistered/burnt | 171 | 135 | **1.61 (1.13-2.29)** | **0.009** |
|  | Mild burn | 161 | 148 | **1.60 (1.13-2.26)** |  |
|  | Tan/No effect | 148 | 207 | **Ref** |  |
| **rs861539 (XRCC3)** | Non-carriers (CC genotype) | 206 | 164 | **Ref** | **0.001** |
|  | Carriers (CT, TT genotype) | 274 | 326 | **0.68 (0.51-0.90)** |  |
| **MC1R<sup>c</sup>** | Non-carriers | 148 | 216 | **Ref** | **0.002** |
|  | Carriers | 332 | 274 | **1.58 (1.18-2.11)** |  |

<sup>a</sup>Odds ratio, OR calculated for each unit decrease in ln(telomere length).
<sup>b</sup>P-values were derived from $\chi^2$ -test, two sided and considered statistically significant if < 0.05, indicated in bold.
<sup>c</sup>The individuals with any one MC1R polymorphism, that include V60L, D84E, V92M, R142H, R151C, I155T, R160W, R163Q, D294H, and T314T, were considered as carriers.

### Mendelian Randomization

Mendelian randomization analysis was carried out to determine the causal effect of telomere length (exposure) on BCC risk (outcome) using single nucleotide polymorphisms (SNPs) as instrumental variables. The SNPs, represented by rs6060627, rs6772228, rs9257445, rs1317082, rs2487999, rs7726159, rs755017, rs412658, rs3027234, associated with telomere length were taken from the previously published genome-wide association studies (30-32).

In a two-sample MR study, the association estimates (β-estimate in terms of kb of telomere length) and the standard errors were determined for the variant allele of each SNP associated with ‘short telomere length’ from the published genome-wide association studies (30-32). The estimates (ln(OR)) for the instrument-outcome association was calculated from logistic regression in this study **(Supplementary material, Supplementary Table S3, Supplementary Table S4)**. Analysis showed no statistically significant association between telomere length and risk of BCC using inverse variance weighted (IVW) method (OR = 0.57, 95%CI = 0.10-3.23, *P* = 0.53) and maximum-likelihood method (OR = 0.57, 95%CI = 0.10-3.25, *P* = 0.52). In addition, the *P* value for heterogeneity test statistic was statistically significant implying failure of instrumental variable assumptions for at least one of the SNPs (*P* = 0.01 with 8 degrees of freedom) **(Supplementary Figure S1 and Supplementary Table S5)**. Since arsenic exposure was found to be a statistically significant confounder, we carried out Mendelian randomization using a one-sample study with non-overlapping datasets to determine the effect of telomere length on BCC risk (33).

For a non-overlapping sample setting, the cases and controls separately were split as two groups each (cases-dataset 1 and 2, controls-dataset 1 and 2) and having similar mean and variance for arsenic exposure. The cases included 261 individuals in both dataset 1 and dataset 2 and the controls 265 individuals in dataset 1 and 264 in dataset 2. The median arsenic exposure within datasets of cases and controls were comparable **(Supplementary Table S6)**. We independently assessed the validity of the instrumental variables assumptions using dataset 1. The variant allele of each SNP was not associated with any of the confounders **(Supplementary material and Supplementary Table S7)**. The estimates of instrument-outcome association were obtained from the logistic regression carried out on dataset 2; the estimates of instrument-exposure association were obtained from the dataset 1 only in the control population **(Supplementary Table S8 and Supplementary Table S9)**. The estimates were adjusted for the measured confounders in the study population that included age, arsenic exposure and country. The combined estimated magnitude of association of all 10 SNPs for a 1 standard deviation decrease in the telomere length was associated with an increased risk of BCC with an OR of 1.83 (95% CI = 0.98-3.40, *P* = 0.06) using the IVW method; OR of 1.83 (95% CI = 0.98-3.41, *P* = 0.06) with the maximum-likelihood method, although not statistically significant. The *P*-value for heterogeneity test statistic for the IVW and maximum-likelihood method was not significant (*P* = 0.10 with 9 degrees of freedom) **(Supplementary Figure S2 and Supplementary Table S10)**.

### Association between arsenic exposure and telomere length

Linear regression model in the combined cases and controls, adjusted for country and age, showed no statistically significant association between decreased telomere length and increase in the arsenic exposure (β = −0.006, 95% CI = −0.02--0.01, *P* = 0.45). However, stratification based on median distribution showed that the association of arsenic exposure on telomere length was statistically significant in the individuals that were exposed to higher than median (>1.32 µg/L) measure of arsenic exposure with a β-estimate -0.026 (95% CI = − 0.05- -0.003, *P* = 0.02) compared to those with lower than median exposure (≤ 1.32 µg/L; β = 0.026, 95% CI = −0.04-0.09, *P* = 0.41). In a similar analysis using WHO’s guideline for arsenic in drinking water (≤ 10µg/L), the estimates were found to be β = −0.043 (95% CI = −0.09- - 0.005, *P* = 0.08) in the higher exposure group (>10 µg/L) and β = 0.011 (95% CI = −0.02-0.04, *P* = 0.45) in the lower exposure group (≤ 10µg/L) **(Table 2)**.

**Table 2.**
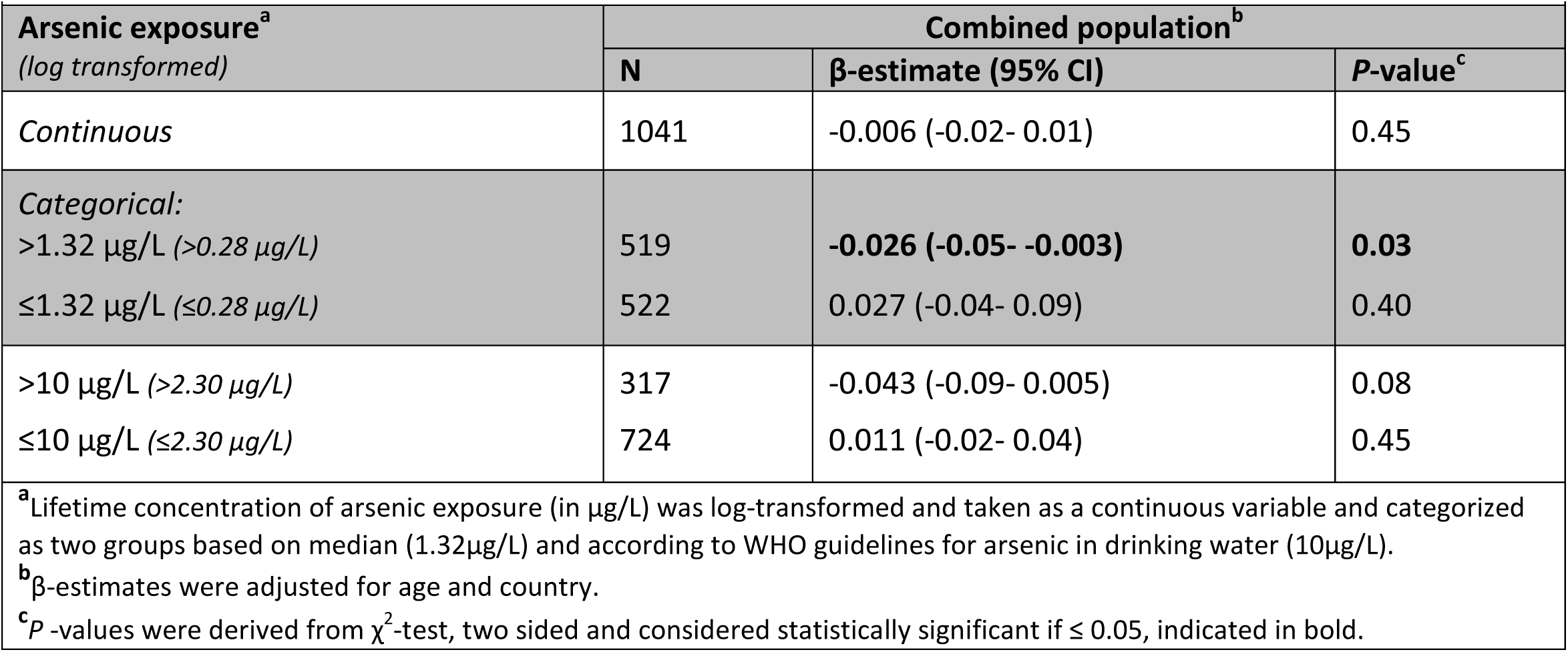
Association between arsenic exposure and telomere length

### Combined effect of arsenic exposure and telomere length on BCC risk

Due to statistically significant interaction (*P* = 0.02) between arsenic exposure and telomere length on the risk of BCC, the data were analyzed for the effect of telomere length on the risk within the three tertiles of arsenic exposure. The individuals with short telomeres in the highest exposure group of arsenic carried statistically significantly increased risk of BCC, q3 (OR = 16.13, 95% CI = 6.71-40.00, *P*<0.0001), followed by those in medium exposure group, q2 (OR = 5.05, 95% CI = 2.29-10.20, *P* <0.0001), and in the low exposure group, q1 (OR = 3.44, 95% CI = 1.81-6.54, *P* = 0.0002). The effect of telomere length on BCC risk within each tertile of arsenic exposure remained statistically significant even after adjustment with different confounders **(Table 3)**.

**Table 3.** Effect of telomere length as a continuous variable on BCC risk in groups with different arsenic exposures

| Arsenic exposure<sup>a</sup><br>(log-transformed) | Telomere length<sup>b</sup><br>(Median, IQR)<br>(log-transformed) | Univariate Analysis |  |  |  | Multivariate Analysis<sup>e</sup> |  |  |  |
| --- | --- | --- | --- | --- | --- | --- | --- | --- | --- |
|  |  | Cases | Controls | OR (95% CI)<sup>c</sup> | P-value<sup>d</sup> | Cases | Controls | OR (95% CI)<sup>c</sup> | P-value<sup>d</sup> |
| q1: ≤0.70 (≤-0.36) | Median: 0.59 (-0.53)<br>IQR: 0.44-0.72 (-0.82- -0.33) | 183 | 163 | **3.44 (1.81-6.54)** | **0.0002** | 166 | 150 | **3.24 (1.54-6.80)** | **0.002** |
| q2: 0.71-16.38 (-0.35- 2.80) | Median: 0.63 (-0.46)<br>IQR: 0.46-0.71 (-0.78- -0.34) | 172 | 175 | **5.05 (2.29-10.20)** | **<0.0001** | 166 | 165 | **2.62 (1.12-6.13)** | **0.03** |
| q3: >16.38 (>2.80) | Median: 0.64 (-0.45)<br>IQR: 0.53-0.74 (-0.63- -0.30) | 163 | 185 | **16.13 (6.71-40.00)** | **<0.0001** | 148 | 175 | **20.00 (6.71-58.82)** | **<0.0001** |

<sup>a</sup>Lifetime concentration of arsenic exposure (in µg/L) was log transformed and grouped into tertiles.
<sup>b</sup>Telomere length was log-transformed and taken as a continuous variable for each group of arsenic exposure.
<sup>c</sup>Odds ratio, OR calculated for each unit decrease in ln(telomere length).
<sup>d</sup>P -values were derived from $\chi^2$ -test, two sided and considered statistically significant if ≤ 0.05, indicated in bold.
<sup>e</sup>Odds ratio in multivariate model was adjusted for age, sex, eye color, country, skin complexion, skin response to sun exposure, *XRCC3* polymorphism, *MC1R* polymorphisms.

We also investigated the combined effect of arsenic exposure and telomere length on BCC risk. The study population was stratified into nine sub-groups based on arsenic exposure and telomere length; three groups of arsenic exposure categorized as high, medium and low exposure and three groups of telomere length categorized as short, medium and long. The group with lowest arsenic exposure and longest telomere length was used as reference and the ORs were determined for the remaining eight groups after adjustment for age and country. The highest risk was observed for the group with shortest telomeres and highest exposure with an OR of 10.56 (95% CI = 5.14-21.70). The interaction between highest arsenic exposure and shortest telomere length was greater than additive (ICR_high_ = 6.56, 95% CI = 0.72-17.90, P<0.03) and multiplicative interaction greater than 1, but not statistically significant (MII_high_ = 1.99, 95% CI = 0.65-5.98, P<0.23). The OR for the risk of BCC in the group with short telomeres and medium exposure was 3.12 (95% CI = 1.73-5.62). Similarly, the group with medium telomeres and high exposure (OR = 2.29, 95% CI = 1.21-4.36) and the group with medium telomeres and medium exposure (OR = 2.25, 95% CI = 1.27-4.00) statistically significantly increased the risk of BCC. However, the additive and multiplicative interactions were not statistically significant in any of the remaining groups **(Figure 3 and Table 4)**.

**Table 4.** Effect of interaction between arsenic exposure and telomere length on risk of BCC

| Telomere length<br>(log-transformed) | Lifetime concentration of arsenic exposure measured in µg/L (log-transformed) |  |  |  |  |  |  |  |  |
| --- | --- | --- | --- | --- | --- | --- | --- | --- | --- |
|  | Lowest exposure<br>Median: 0.61 (-0.49)<br>IQR: 0.48-0.70 (-0.73- -0.36) |  |  | Medium exposure<br>Median: 1.32 (0.28)<br>IQR: 0.93-2.75 (-0.07- 1.01) |  |  | Highest exposure<br>Median: 26.02 (3.26)<br>IQR: 16.38-41.90 (2.80- 3.74) |  |  |
|  | Cases | Controls | OR (95%CI)<sup>a</sup> | Cases | Controls | OR (95%CI)<sup>a</sup> | Cases | Controls | OR (95%CI)<sup>a</sup> |
| **Longest**<br>Median: 0.78 (-0.25)<br>IQR: 0.73-0.89 (-0.31- -0.12) | 72 | 21 | Reference | 50 | 72 | 0.99 (0.56-1.75) | 41 | 92 | 1.76 (0.93-3.35) |
| **Medium**<br>Median: 0.61 (-0.49)<br>IQR: 0.57-0.65 (-0.56- -0.43) | 71 | 36 | **2.15 (1.21-3.82)** | 60 | 51 | **2.25 (1.27-4.00)** | 41 | 88 | **2.29 (1.21-4.36)** |
| **Interaction with arsenic exposure<sup>b</sup>**<br>MII (95% CI)<br>ICR (95% CI) |  |  |  | 1.06 (0.38-2.76) *P* < 0.90<br>0.08 (-2.04-2.11) *P* < 0.93 |  |  | 0.60 (0.21-1.76) *P* < 0.37<br>-0.69 (-3.10-1.72) *P* < 0.54 |  |  |
| **Shortest**<br>Median: 0.43 (-0.84)<br>IQR: 0.35-0.49 (-1.05- -0.71) | 90 | 47 | **3.05 (1.76-5.28)** | 57 | 45 | **3.12 (1.73-5.62)** | 36 | 71 | **10.56 (5.14-21.70)** |
| **Interaction with arsenic exposure<sup>b</sup>**<br>MII (95% CI)<br>ICR (95% CI) |  |  |  | 1.04 (0.38-2.76) *P* < 0.93<br>0.05 (-2.66-2.84) *P* < 0.98 |  |  | **1.99 (0.65-5.98) *P* < 0.23**<br>**6.56 (0.72-17.90) *P* < 0.03** |  |  |

<sup>a</sup>Odds ratio, OR and 95% confidence interval (CI) for the risk associated with combined effect of arsenic exposure and telomere length on BCC risk were adjusted for age and country. Global *P*-value for the combined effect was <0.0001.
<sup>b</sup>MI is multiplicative interaction index and ICR, interaction constant ratio. The lowest arsenic exposure and longest telomeres was used as a single reference group for the interaction analysis.

## Discussion

In the present multi-center case-control study, we observed that in contrast to earlier reports, the skin BCC patients carried statistically significantly shorter telomeres than the healthy controls (11). However, the study population included individuals exposed to arsenic through drinking water, which has been shown to affect both BCC risk and telomere length. We observed that higher than the median exposure to arsenic (1.32µg/L) associated with statistically significant decrease in telomere length. The analysis of the effect of telomere length, within the tertiles based on arsenic exposure, showed that the individuals with shorter telomeres in the highest exposure groups were at the largest risk of BCC followed by those in the low and medium exposure groups. In a greater than additive interaction we observed more than 10-fold increased risk of BCC in the sub-group with the highest arsenic exposure and shortest telomeres compared to the group with lowest exposure and longest telomeres.

The observational studies have consistently shown statistically significant associations between long telomeres and increased cancer risk (27,35,36). The genetic basis for those observations are provided by the studies showing that various polymorphisms that modulate telomere length also affect the risk of different cancers, with alleles segregating with increased telomere length associating with increased risk (10,11,37). Long telomeres are postulated to increase the proliferative potential by delaying senescence (38). A Mendelian randomization study based on 3361 BCC cases and 11518 controls from the skin BCC case-control study nested within Nurse’s Health study (NHS) and Health Professionals Follow-up study (HPFS), reported an increased cancer risk with genetically increased telomere length (11). Contrary to those reported observations, our data analysis showed that decreased telomere length associated with an increased BCC risk due to confounding effect of arsenic exposure. However, it may be pointed out that several telomeropathies, besides various debilitating disorders also predispose individuals to some cancers including acute myeloid leukemia and tongue carcinoma (13,39). The telomeropathies-caused by inherited mutations in genes involved in telomere structure maintenance, repair, replication and preservation of equilibrium-result in critically short telomeres (7,13,40).

The lack of genetic basis for our observation of association between short telomeres and increased BCC risk was confirmed by the failure of Mendelian randomization using a two-sample study. A statistically significant heterogeneity in the two-sample model indicated that the subjects used to estimate the SNP-outcome and SNP-exposure associations are not homogeneous, due to confounding in the exposure-outcome relationship, arsenic exposure in this case (41). That was further corroborated in one sample study that incorporated arsenic exposure into the model and showed association between decreased telomere length and increased BCC risk (33). However, one-sample Mendelian randomization study is generally limited due to weak instrument bias and underestimation of true causal effect (41,42).

Thus, the analysis of our data indicated that the observed direction of the effect of telomere length on BCC risk was mainly due to arsenic exposure through drinking water. The relationship between arsenic exposure and telomere length has been assessed in both experimental and observational studies (12,43,44). One of the principle mechanisms of arsenic toxicity is oxidative stress. Experimental studies have shown that arsenic-induced oxidative stress leads to telomere attrition, DNA damage, chromosomal instability and apoptosis (45,46). *In vitro* studies on arsenic exposure suggested that the lower concentrations of arsenic exposure (<1µM) either maintain or increase the telomere length, whereas, the higher concentrations (>1µM) result in a drastic decrease in the telomere length (44,46). Due to the interaction between arsenic exposure and telomere on skin lesions, different magnitudes of the effect dependent on levels arsenic have been predicted (43). The direction of the effects of the exposure on telomere length, besides the arsenic dose, has also been reported to be dependent on factors such as duration of exposure, age and DNA repair capacity (21,47). Studies have indicated that arsenic tends to accumulate in the skin and associates with BCC and other non-melanoma skin cancers (48).

The decreased telomere length through arsenic exposure being causal for the increase in BCC risk, as observed in this study, cannot be ruled out; however, that contrasts with epidemiological studies that have consistently associated longer telomere with increased cancer risk including BCC (11). We, therefore, hypothesize that probably the observed effect on BCC risk is driven through a combination of arsenic exposure and telomere shortening, which is supported by greater than additive interaction implying a causal interaction between the two factors (49). Previously, epidemiological studies have shown apparent synergistic interactions between arsenic and other risk factors of skin lesions such as smoking, fertilizer use and sunlight exposure (50,51). Our study is the first to show a synergistic effect of arsenic exposure and reduced telomere length on the risk of BCC.

In summary, in contrast to consistent reports of association between increased telomere length and increased risk of various cancers, our data showed that in a population exposed to arsenic, it is the short telomeres that associated with increased BCC risk (11). The results of the data analysis showed that the arsenic exposure modulates the direction of the effect of telomere length. We also observed a synergistic interaction between arsenic exposure and telomere length in the group with the highest exposure and shortest telomeres.

## Supporting information

