## Supplementary material for "Telomere length, arsenic exposure and risk of basal cell carcinoma of skin"

**Running title:** Telomere length and basal cell carcinoma risk

***Corresponding author:** Rajiv Kumar, Division of Molecular Genetic Epidemiology, German Cancer Research Center, Im Neunheimer Feld 580, 69120 Heidelberg, Germany

**Key words:** telomere length; arsenic exposure; basal cell carcinoma.

### Supplementary Materials and Methods

#### Association of co-variates and BCC risk

The associations between BCC risk and the co-variates such as age (continuous variable), sex (male or female), eye color (blue/ green, hazel/ brown and others), skin complexion (light, medium and dark), skin response to sun exposure (blistered/burnt, mild burn and tan), body mass index (continuous), were estimated using univariate logistic regression. The corresponding odds ratios (OR) and 95% confidence intervals (CI) were also determined. The arsenic exposure values measured as time weighted average concentration in µg/l over the lifetime of an individual were log-transformed. Arsenic exposure as a categorical variable was used based on tertile distribution (q1 = ≤0.70, q2 = 0.71-16.38, q3 >16.38) and was adjusted for country.

#### Genotyping

The DNA from the BCC Cases and controls were genotyped for nine single nucleotide polymorphisms (SNPs) that are chosen from previously published genome-wide association studies, which showed significant association with telomere length [1-3]. The polymorphisms and genes investigated included: rs6060627 (*BCL2L1*), rs6772228 (*PXK*), rs9257445 (*ZNF311*), rs1317082 (*TERC*), rs2487999 (*OBFC1*), rs7726159 (*TERT*), rs755017 (*RTEL1*), rs412658 (*ZNF676*), rs3027234 (*CTC1*). The *MC1R* polymorphisms (V60L, D84E, V92M, R142H, R151C, I155T, R160W, R163Q, D294H and T314T) and *XRCC3* polymorphism rs861539, genotyped previously, were included in the analysis [4,5]. The genotyping was performed using TaqMan® SNP Genotyping assays. PCR was performed in a 5-10µl reaction volume in an optical 384-well reaction plate and consisted of 5µl of 5-10ng/µl DNA as template, 1µl 5X HOT FIREPol probe qPCR Mix Plus with ROX (Solis BioDyne, Tartu, Estonia) and 1X probe-primer mix. The initial temperature condition for PCR was set at 95$^{o}$C/15 min followed by 35-

40 cycles at 94$^{o}$C/20s and 60$^{o}$C/1 min. Post amplification products were analyzed on a ViiA-7 real-time PCR system (Applied Biosystems) and the fluorescence intensities of each sample were measured. The genotypes were determined visually based on the fluorescence emission data depicted on the X-Y scatter plot of the ViiA7 RUO software, version 1.2.2 (Applied Biosystems). Genotype frequencies of all SNPs were tested in control subjects for deviation from Hardy-Weinberg equilibrium using Pearson's $\chi^2$ test.

**Assessment of SNPs as valid instrumental variables for one-sample MR study**

The polymorphisms were considered as instrumental variables based on the following assumptions: i) the polymorphisms are associated with telomere length. ii) The polymorphisms are independent of any confounders of the telomere length-BCC risk association iii) the polymorphisms are independent of the outcome given the exposure and confounders. All the nine SNPs associated with telomere length were selected based on statistical significance from genome-wide association studies [1-3]. Each SNP represented different genes and were present on different chromosomes, therefore not genetically linked. Linear regression model was computed to determine the SNP-telomere length association using telomere length as dependent variable with polymorphism, age (continuous), arsenic exposure (continuous), country and polymorphism*age as independent variables using controls from dataset 1. F-value was obtained from linear regression analysis; an F-value above 10 indicates sufficient statistical strength of the polymorphisms to be used as valid instruments [6,7]. Pleiotropy was assessed using logistic regression analysis based on the allele status for each SNP and each of the significant confounders (age, sex, body mass index, eye color, skin complexion and skin response to sun exposure) for only control population. Linear and logistic regression models were adjusted for the confounding effect of arsenic in the data.

### Supplementary Results

#### Association of co-variates on risk of BCC

Other co-variates that showed statistically significant association with increased risk of BCC included age at diagnosis (continuous variable, $P$<0.0001), sex (females versus males, $P$ = 0.03), eye color (Hazel/brown versus Blue/ green or others, $P$ = 0.07), skin complexion (light versus medium/dark complexion, $P$<0.0001), skin response to sun exposure (blistered/burnt or mild versus tan/no effect, $P$ = 0.0005). Body mass index as a continuous variable did not show any statistically significant association with risk of BCC ($P$ = 0.30). In addition, the risk association for individuals with, compared to those without *MC1R* genetic variants; and in carriers of rs861539 *(XRCC3)* polymorphism compared to the non-carriers, was previously reported on the present study population and the data was included in the analysis [4,5,8]. Stratification of arsenic exposure based on tertile distribution, adjusted for country, showed that the individuals in higher exposure group of arsenic exposure, q3 (>16.38) compared to those in lower exposure group, q1 (≤0.70) were at statistically significantly increased risk of BCC (OR = 1.83, 95%CI = 1.18-2.83), and the individuals in the medium exposure group were not associated with the risk of BCC (OR = 0.97, 95% CI = 0.71-1.34) **(Supplementary Table S2)**.

#### SNP-BCC risk and SNP-Telomere length estimates for two-sample MR study

The estimates for SNP-BCC risk association were determined using logistic regression analysis. Analysis of genotype data showed that out of the nine SNPs, the carriers of the variant allele (C-allele) for rs9257445, located within the *ZNF311* locus, were at statistically significantly decreased the risk of BCC (OR = 0.68, 95% CI = 0.56-0.83, *P* = 0.0002). However, in the data from the genome-wide association study, the variant allele for the rs9257445 polymorphism was shown to be associated with decreased telomere length [1]. The estimates for BCC risk for the variant allele for rs9257445 was in the opposite direction. The analysis of the genotyping data of the remaining polymorphisms showed that the carriers of the variant alleles for the rs6772228 (*PXK*), rs1317082 (*TERC*), rs7726159 (*TERT*) and rs3027234 (*CTC1*) showed a decreased risk for BCC and the carriers of the variant alleles for the rs6060627 (*BCL2L1*), rs2487999 (*OBFC1*), rs755017 (*RTEL1*) and rs412658 (*ZNF676*) showed an increased risk of BCC, however the associations were not statistically significant **(Supplementary Table S3)**. The estimates for the SNP-telomere length association were determined from genome-wide association studies **(Supplementary Table S4)** [1-3].

**Polymorphisms as valid instrumental variables for one sample MR study**

A one-sample MR study was conducting using non-overlapping datasets from the same population. The cases and controls separately were split as two groups each (cases-dataset 1 and 2, controls-dataset 1 and 2) and having similar mean and variance for arsenic exposure. The assumptions for the SNPs to be considered as valid instrumental variables were assessed independently.

Assumption 1: Linear regression, adjusted for age, country and arsenic exposure, was carried out only in the control population from the dataset 1. The genetic associations obtained could be biased when using exposure estimates in the combined population and hence only the estimates from the controls were used [9]. No significant association between any of the SNPs and telomere length was observed in our data. The β-estimates for three SNPs, rs9257445, rs755017 and rs412568 deviated from the estimates of telomere length reported in the genome-wide association studies [1-3]. However, F-statistic >10 was observed for each individual SNP, therefore the SNPs were considered as valid instruments to be used in MR analysis [7] **(Supplementary Table S9)**.

Assumption 2: The regression analysis showed no association between the variant allele of each polymorphism and any of the other confounders in the study (age, sex, body mass index, eye color, skin complexion and skin response to sun exposure) indicating no pleiotropy in the study population **(Supplementary Table S7)**.

Assumption 3: Logistic regression analysis was carried out in the dataset 2, adjusted for age, country and arsenic exposure. Analysis of genotyping data showed that out of the nine SNPs, the carriers of the variant allele (C-allele) for rs9257445, located within the *ZNF311* locus, were at statistically significantly decreased risk of BCC (OR = 0.73, 95% CI = 0.54-0.99, $P$ = 0.04). The variant allele for rs9257445 in our study population was associated with longer telomeres and hence the estimates for BCC risk showing decreased risk was in the expected direction. The analysis of the genotyping data of the remaining polymorphisms showed that the carriers of the variant alleles for rs6772228 (*PXK*), rs7726159 (*TERT*) and rs412658 (*ZNF676*) and rs3027234 (*CTC1*) showed a decreased risk for BCC and the carriers of the variant alleles for rs1317082 (*TERC*), rs2487999 (*OBFC1*), rs755017 (*RTEL1*) showed an increased risk of BCC, however the associations were not statistically significant. The variant allele for the rs6060627 (*BCL2L1*) polymorphism was not associated with the risk. The *XRCC3* polymorphism (rs861539) was also included in the Mendelian randomization analysis. Although, there is no evidence for the association between the polymorphism in *XRCC3* and telomere length, the carrier of the variant allele (T-allele) for rs861539 showed a statistically significant decreased risk of BCC in our study (OR = 0.69, 95% CI = 0.52- 0.91, $P$ = 0.009) and hence was included in the analysis **(Supplementary Table S8)**.

**Legends to supplementary figures**

**Supplementary Figure S1:** Two-sample Mendelian randomization showing association between telomere-length associated polymorphisms and risk of BCC. The scatter plots show the per-allele association with BCC risk plotted on the y-axes (represented as natural log of odds ratio) against the per-allele association with kb of telomere length on x-axes from genome-wide association studies. The continuous blue line represents the Mendelian randomization estimate of telomere length on BCC risk. The model included all the nine SNPs chosen from genome-wide association studies. The point estimate using the inverse-variance weighted method along with 95% confidence intervals are shown above the scatter plot.

**Supplementary Figure S2:** One-sample Mendelian randomization using non-overlapping datasets showing association between telomere-length associated polymorphisms and risk of BCC. The scatter plots show the per-allele association with BCC risk plotted on the y-axes (represented as natural log of odds ratio) against the per-allele association with telomere length on x-axes from this study. The continuous blue line represents the Mendelian randomization estimate of telomere length on BCC risk. The model included all the nine SNPs chosen from genome-wide association studies and rs861539 (*XRCC3* polymorphism). The point estimate using the inverse-variance weighted method along with 95% confidence intervals are shown above the scatter plot.

**Supplementary Figure S1**

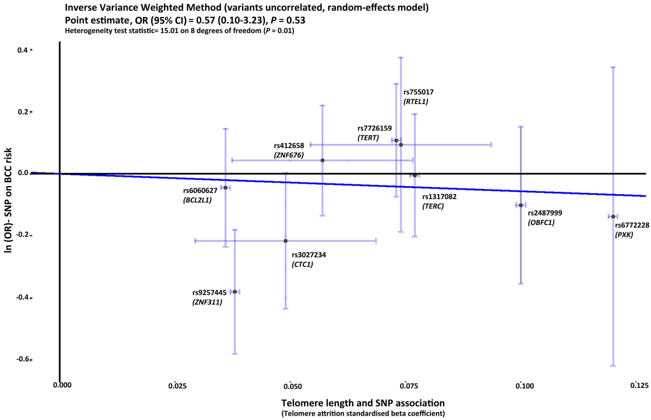

**Supplementary figure S2**

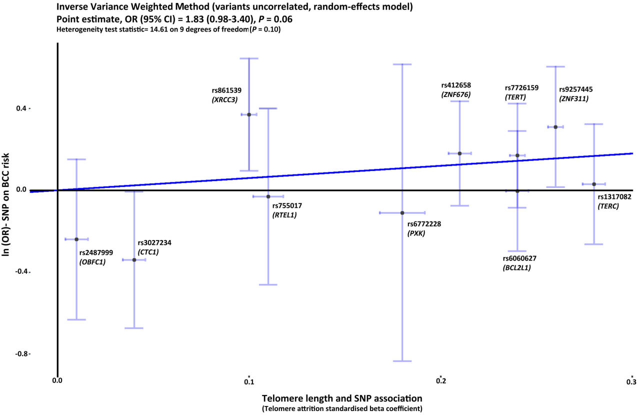

**Supplementary Table S1.** Primers for telomere length measurement

| | Size (bp) | 5'→ 3' Sequence | Ta ($^oC$) |
|---|---|---|---|
| **Telomeres** | | | |
| telg | 79 | 5'ACACTAAGGTTTGGGTTTGGGTTTGGGTTTGGGTTAGTGT3' | 59 $^oC$ |
| telc | | 5' TGTTAGGTATCCCTATCCCTATCCCTATCCCTATCCCTAACA3' | |
| **Albumin** | | | |
| albugcr2 | 98 | 5'CGGCGGCGGGCGGCGCGGGCTGGGCGGCCATGCTTTTCAGC TCTGCAAGTC3' | 85 $^oC$ |
| albdgcr2 | | 5'GCCCGGCCCGCCGCGCCCGTCCCGCCGAGCATTAAGCTCTTT GGCAACGTAGGTTTC3' | |

**Supplementary Table S2.** Univariate Analysis of BCC risk due to different factors

| Patient Characteristics | | Cases N=528(%) | Controls N=533(%) | OR (95% CI) | *P*-value[c] |
|---|---|---|---|---|---|
| **Telomere length[a]** *(log-transformed)* | *Continuous* | 524 (99.2) | 527 (98.9) | **5.92 (3.92-9.01)** | **<0.0001** |
| | Median: 0.62 *(-0.48)* | | | | |
| | IQR: 0.49-0.73 *(-0.71- -0.31)* | | | | |
| | *Categorical* | | | | |
| | q1: ≤0.49 *(-0.71)* | 237 (44.9) | 105 (19.7) | **4.74 (3.46-6.50)** | |
| | q2: 0.50-0.73 *(-0.70- -0.31)* | 167 (31.6) | 170 (31.9) | **2.06 (1.52-2.80)** | |
| | q3: >0.73 *(-0.31)* | 120 (22.7) | 252 (47.3) | **Ref** | **<0.0001** |
| | Missing | 4 (0.8) | 6 (1.1) | | |
| **Age** | *Continuous* | 528 (100) | 533 (100) | **1.04 (1.03-1.05)** | **<0.0001** |
| | Median (IQR): 64 (55-72) years | | | | |
| **Sex** | Male | 236 (44.7) | 274 (51.4) | **Ref** | |
| | Female | 292 (55.3) | 259 (48.6) | **1.31 (1.03-1.67)** | **0.03** |
| **Body Mass Index (in Kg/m[3])** | *Continuous* | 500 (94.7) | 510 (95.7) | 0.99 (0.96-1.01) | 0.30 |
| | Median (IQR) = 26.55 (23.88-29.39) | | | | |
| | missing | 28 (5.3) | 23 (4.3) | | |
| **Eye color** | Blue/Green | 301(57.0) | 270 (50.7) | **1.36 (1.05-1.77)** | |
| | Hazel/Brown | 171 (32.4) | 209 (39.2) | **Ref** | 0.07 |
| | Others | 28 (5.3) | 30 (5.6) | 1.14 (0.66-1.98) | |
| | missing | 28 (5.3) | 24 (4.5) | | |
| **Arsenic exposure[b]** *(log-transformed)* | q1: ≤0.70 *(≤-0.36)* | 184 (34.9) | 164 (30.8) | **Ref** | **0.006** |
| | q2: 0.71-16.38 *(-0.35- 2.80)* | 174 (33.0) | 179 (33.6) | 0.97 (0.71-1.34) | |
| | q3: >16.38 *(>2.80)* | 164 (31.1) | 186 (34.9) | **1.83 (1.18-2.83)** | |
| | missing | 6 (1.0) | 4 (0.7) | | |
| **Skin Complexion** | light | 280 (53.0) | 212 (39.8) | **1.71 (1.34-2.18)** | **<0.0001** |
| | medium/dark | 247 (46.8) | 320 (60.0) | **Ref** | |
| | missing | 1 (0.2) | 1 (0.2) | | |
| **Skin response to sun exposure** | blistered/burnt | 185 (35.0) | 141 (26.5) | **1.80 (1.33-2.42)** | |
| | mild burn | 169 (32.0) | 160 (30.0) | **1.45 (1.08-1.94)** | |
| | tan | 163 (30.9) | 223 (41.8) | **Ref** | **0.0005** |
| | missing | 11 (2.1) | 9 (1.7) | | |

[a]Odds ratio, OR calculated for each unit decrease in ln(telomere length).
[b]Lifetime concentration of arsenic exposure (in µg/L) was log-transformed and categorized as tertiles, Odds Ratio, OR adjusted for Country.
[c]*P* -values were derived from $\chi^2$-test, two sided and considered statistically significant if < 0.05, indicated in bold.

**Supplementary table S3**. Telomere length associated SNPs and risk of BCC

| **SNPs associated with Telomere Length** | | **Cases (N=1056)** | **Controls (N=1066)** | **OR (95% CI)[a]** | ***P*-value[b]** |
|---|---|---|---|---|---|
| **rs6060627** (***BCL2L1***) | Major allele (C-allele)<br>Minor allele (T-allele)<br>missing | 753<br>301<br>2 | 764<br>292<br>10 | 1.05 (0.87-1.27) | 0.64 |
| **rs6772228** (***PXK***) | Major allele (T-allele)<br>Minor allele (A-allele)<br>missing | 1020<br>32<br>4 | 1027<br>37<br>2 | 0.87 (0.54-1.41) | 0.57 |
| **rs9257445** (***ZNF311***) | Major allele (G-allele)<br>Minor allele (C-allele)<br>missing | 807<br>223<br>26 | 752<br>304<br>10 | **0.68 (0.56-0.83)** | **0.0002** |
| **rs1317082** (***TERC***) | Major allele (A-allele)<br>Minor allele (G-allele)<br>missing | 778<br>262<br>16 | 786<br>266<br>14 | 0.99 (0.81-1.20) | 0.88 |
| **rs2487999** (***OBFC1***) | Major allele (C-allele)<br>Minor allele (T-allele)<br>missing | 857<br>141<br>58 | 921<br>137<br>8 | 1.11 (0.86-1.42) | 0.43 |
| **rs7726159** (***TERT***) | Major allele (C-allele)<br>Minor allele (A-allele)<br>missing | 715<br>339<br>2 | 691<br>365<br>10 | 0.90 (0.75-1.08) | 0.24 |
| **rs755017** (***RTEL1)*** | Major allele (A-allele)<br>Minor allele (G-allele)<br>missing | 920<br>112<br>24 | 956<br>106<br>4 | 1.10 (0.83-1.45) | 0.51 |
| **rs412658** (***ZNF676)*** | Major allele (C-allele)<br>Minor allele (T-allele)<br>missing | 645<br>397<br>14 | 653<br>385<br>28 | 1.04 (0.87-1.25) | 0.63 |
| **rs3027234** (***CTC1)*** | Major allele (C-allele)<br>Minor allele (T-allele)<br>missing | 871<br>185<br>0 | 815<br>215<br>36 | 0.81 (0.65-1.00) | 0.05 |

[a]Odds ratio, OR and 95% confidence intervals (CI) was calculated in 528 patients and 533 healthy controls.
[b]*P* -values were derived from $\chi^2$-test, two sided and considered statistically significant if < 0.05, indicated in bold.

**Supplementary Table S4.** Mendelian randomization estimates for two-sample study

| SNP | Chr. | Position (hg19) | Gene | Allele (Major/minor) | MAF | Instrument-exposure association from genome-wide association studies | | Instrument-outcome association from present study | | Ref |
|---|---|---|---|---|---|---|---|---|---|---|
| | | | | | | β-estimate[a] | Standard error | Estimate[b] | Standard error | |
| rs6060627 | 20 | 30262159 | *BCL2L1* | C/T | 0.28 | 0.036 | 0.0005 | 0.0449 | 0.0969 | [1] |
| rs6772228 | 3 | 58376019 | *PXK* | T/A | 0.04 | -0.12 | 0.0005 | -0.1381 | 0.2454 | [1] |
| rs9257445 | 6 | 28949206 | *ZNF311* | G/C | 0.29 | -0.038 | 0.0005 | -0.3804 | 0.1017 | [1] |
| rs1317082 | 3 | 169497585 | *TERC* | A/G | 0.26 | -0.077 | 0.0005 | -0.00492 | 0.1007 | [1] |
| rs2487999 | 10 | 105659826 | *OBFC1* | C/T | 0.13 | 0.1 | 0.0005 | 0.1009 | 0.129 | [1] |
| rs7726159 | 5 | 1282319 | *TERT* | C/A | 0.35 | 0.073 | 0.0005 | -0.108 | 0.0924 | [1] |
| rs755017 | 20 | 62421622 | *RTEL1* | A/G | 0.10 | -0.074 | 0.01 | 0.0934 | 0.1432 | [2] |
| rs412658 | 19 | 22359440 | *ZNF676* | C/T | 0.36 | -0.057 | 0.01 | 0.043 | 0.0905 | [3] |
| rs3027234 | 17 | 8136092 | *CTC1* | C/T | 0.20 | -0.049 | 0.01 | -0.2165 | 0.1115 | [3] |

[a]The beta-estimates for the variant allele of each SNP associated with telomere length, determined from genome-wide association studies.
[b]The effect estimates were determined using logistic regression.

**Supplementary Table S5.** Mendelian randomization estimates for nine genome-wide association studies based SNPs with BCC risk in two sample study

| **Method** | **Estimate (95% CI)[a]** | **SE** | **OR (95% CI)[b]** | ***P*-value** | **Heterogeneity test statistic (*P*-value)** |
|---|---|---|---|---|---|
| ***Inverse-Variance Weighted*** | -0.56 (-2.29 to 1.17) | 0.89 | 0.57 (0.10-3.23) | 0.53 | **0.01** |
| ***Maximum-Likelihood*** | -0.57 (-2.31 to 1.18) | 0.92 | 0.57 (0.10-3.25) | 0.52 | |

[a]Mendelian randomization estimates for short telomeres on BCC risk.
[b]Odds Ratio, OR and 95% confidence interval, CI were calculated by taking the exponential of the estimates. SE, standard error.

**Supplementary Table S6.** Non-overlapping datasets of cases and controls for a one-sample Mendelian randomization study based on arsenic exposure

| Arsenic exposure | Controls | |
|---|---|---|
| | Dataset 1 | Dataset 2 |
| N | 265 | 264 |
| Mean | 12.29 | 12.29 |
| Median | 1.86 | 1.81 |
| Standard Deviation | 20.85 | 20.84 |
| Variance | 434.67 | 434.64 |
| **Arsenic exposure** | **Cases** | |
| | **Dataset 1** | **Dataset 2** |
| N | 261 | 261 |
| Mean | 12.94 | 12.97 |
| Median | 1.14 | 1.02 |
| Standard Deviation | 23.28 | 23.32 |
| Variance | 541.90 | 543.77 |

| **Supplementary Table S7.** Association of covariates and telomere-associated SNPs in controls (dataset1) | | | | | | | | | | |
|---|---|---|---|---|---|---|---|---|---|---|
| | | **rs6060627** | | | **rs6772228** | | | **rs9257445** | | |
| **Study-specific co-variates: Patient characteristics** | | **Major Allele (C)** | **Minor Allele (T)** | **OR (95% CI)[a]** | **Major Allele (T)** | **Minor Allele (A)** | **OR (95% CI)[a]** | **Major Allele (G)** | **Minor Allele (C)** | **OR (95% CI)[a]** |
| **Age** | ***Continuous*** | 376 | 158 | 1.00 (0.98-1.01) | 515 | 21 | 0.99 (0.95-1.03) | 375 | 159 | 1.00 (0.99-1.02) |
| **Sex** | **Male**<br>**Female** | 197<br>77 | 181<br>81 | **Ref**<br>1.15 (0.79-1.66) | 263<br>252 | 9<br>12 | **Ref**<br>1.39 (0.58-3.36) | 183<br>192 | 87<br>72 | **Ref**<br>0.79 (0.54-1.14) |
| **Body mass index** | ***Continuous*** | 357 | 151 | 0.97 (0.94-1.01) | 489 | 21 | 0.95 (0.86-1.04) | 353 | 153 | 1.01 (0.97-1.05) |
| **Eye color** | **Blue/Green**<br>**Hazel/Brown**<br>**Others** | 188<br>148<br>21 | 84<br>58<br>9 | 1.14 (0.77-1.70)<br>**Ref**<br>1.09 (0.47-2.53) | 261<br>198<br>30 | 11<br>8<br>2 | 1.04 (0.41-2.64)<br>**Ref**<br>1.65 (0.33-8.14) | 184<br>142<br>27 | 88<br>60<br>5 | 1.13 (0.76-1.68)<br>**Ref**<br>0.44 (0.16-1.19) |
| **Arsenic exposure[b]** *(log transformed)* | ***Continuous*** | 371 | 157 | 1.02 (0.89-1.18) | 508 | 20 | 1.15 (0.83-1.59) | 370 | 156 | 0.93 (0.81-1.06) |
| **Skin Complexion** | **Light**<br>**Medium/Dark** | 155<br>223 | 63<br>95 | 0.95 (0.65-1.39)<br>**Ref** | 207<br>11 | 308<br>10 | 1.64 (0.68-3.92)<br>**Ref** | 154<br>221 | 64<br>95 | 0.97 (0.66-1.41)<br>**Ref** |
| **Skin response to sun exposure** | **Burnt/Blistered**<br>**Mild burn**<br>**Tan** | 105<br>130<br>135 | 41<br>44<br>71 | 0.74 (0.47-1.18)<br>0.64 (0.41-1.01)<br>**Ref** | 136<br>170<br>199 | 8<br>4<br>9 | 1.30 (0.49-3.46)<br>0.52 (0.16-1.72)<br>**Ref** | 102<br>120<br>146 | 44<br>52<br>60 | 1.05 (0.66-1.67)<br>1.05 (0.68-1.64)<br>**Ref** |

[a]Odds ratio, OR calculated for minor allele of each SNP in controls.
[b]Arsenic exposure adjusted for country.

**Supplementary Table S7 (continued).** Association of covariates and telomere-associated SNPs in controls (dataset1)

| Study-specific co-variates: Patient characteristics | | rs1317082 | | | rs2487999 | | | rs7726159 | | |
|---|---|---|---|---|---|---|---|---|---|---|
| | | **Major Allele (A)** | **Minor Allele (G)** | **OR (95% CI)[a]** | **Major Allele (C)** | **Minor Allele (T)** | **OR (95% CI)[a]** | **Major Allele (C)** | **Minor Allele (A)** | **OR (95% CI)[a]** |
| **Age** | ***Continuous*** | 393 | 137 | 0.98 (0.97-1.01) | 456 | 80 | 0.99 (0.97-1.01) | 352 | 184 | 0.99 (0.98-1.01) |
| **Sex** | **Male** | 202 | 66 | **Ref** | 230 | 44 | **Ref** | 186 | 88 | **Ref** |
| | **Female** | 191 | 71 | 1.14 (0.77-1.68) | 226 | 36 | 0.83 (0.52-1.34) | 166 | 96 | 1.22 (0.86-1.75) |
| **Body mass index** | ***Continuous*** | 372 | 130 | 0.98 (0.94-1.02) | 431 | 77 | 1.02 (0.97-1.07) | 334 | 174 | 0.99 (0.96-1.03) |
| **Eye color** | **Blue/Green** | 207 | 63 | 0.77 (0.51-1.17) | 233 | 37 | 0.75 (0.46-1.24) | 178 | 92 | 0.90 (0.62-1.32) |
| | **Hazel/Brown** | 145 | 57 | **Ref** | 170 | 36 | **Ref** | 131 | 75 | **Ref** |
| | **Others** | 19 | 11 | 1.47 (0.66-3.29) | 27 | 5 | 0.87 (0.32-2.42) | 25 | 7 | 0.49 (0.20-1.19) |
| **Arsenic exposure[b]** *(log transformed)* | ***Continuous*** | 386 | 136 | 0.88 (0.76-1.02) | 451 | 77 | 1.01 (0.85-1.21) | 346 | 182 | 1.05 (0.92-1.20) |
| **Skin Complexion** | **Light** | 160 | 54 | 0.95 (0.64-1.41) | 190 | 30 | 0.84 (0.52-1.37) | 141 | 79 | 1.13 (0.78-1.62) |
| | **Medium/Dark** | 233 | 83 | **Ref** | 266 | 50 | **Ref** | 211 | 105 | **Ref** |
| **Skin response to sun exposure** | **Burnt/Blistered** | 111 | 35 | 0.83 (0.51-1.36) | 117 | 29 | 1.64 (0.93-2.92) | 85 | 61 | 1.39 (0.90-2.16) |
| | **Mild burn** | 126 | 44 | 0.92 (0.58-1.46) | 152 | 22 | 0.96 (0.53-1.75) | 124 | 50 | 0.78 (0.51-1.21) |
| | **Tan** | 148 | 56 | **Ref** | 179 | 27 | **Ref** | 136 | 70 | **Ref** |

[a]Odds ratio, OR calculated for minor allele of each SNP in controls.
[b]Arsenic exposure adjusted for country.

**Supplementary Table S7 (continued).** Association of covariates and telomere-associated SNPs in controls (dataset1)

| Study-specific co-variates: Patient characteristics | | rs755017 | | | rs412658 | | | rs3027234 | | |
|---|---|---|---|---|---|---|---|---|---|---|
| | | **Major Allele (A)** | **Minor Allele (G)** | **OR (95% CI)[a]** | **Major Allele (C)** | **Minor Allele (T)** | **OR (95% CI)[a]** | **Major Allele (C)** | **Minor Allele (T)** | **OR (95% CI)[a]** |
| **Age** | ***Continuous*** | 479 | 57 | 1.00 (0.98-1.03) | 344 | 184 | 1.01 (0.99-1.02) | 409 | 111 | 0.99 (0.98-1.01) |
| **Sex** | **Male**<br>**Female** | 237<br>242 | 35<br>22 | **Ref**<br>0.62 (0.35-1.08) | 172<br>172 | 96<br>88 | **Ref**<br>0.92 (0.64-1.31) | 213<br>196 | 53<br>58 | **Ref**<br>1.19 (0.78-1.81) |
| **Body mass index** | ***Continuous*** | 453 | 55 | 0.99 (0.94-1.05) | 322 | 178 | 0.98 (0.95-1.02) | 388 | 104 | 1.01 (0.96-1.05) |
| **Eye color** | **Blue/Green**<br>**Hazel/Brown**<br>**Others** | 246<br>178<br>30 | 24<br>28<br>2 | 0.62 (0.35-1.11)<br>**Ref**<br>0.42 (0.10-1.87) | 170<br>129<br>23 | 92<br>77<br>9 | 0.91 (0.62-1.33)<br>**Ref**<br>0.66 (0.29-1.49) | 205<br>157<br>26 | 55<br>43<br>6 | 0.98 (0.63-1.54)<br>**Ref**<br>0.84 (0.33-2.18) |
| **Arsenic exposure[b]** *(log transformed)* | ***Continuous*** | 471 | 57 | 1.20 (0.98-1.47) | 340 | 182 | 0.94 (0.82-1.08) | 403 | 109 | 0.89 (0.76-1.04) |
| **Skin Complexion** | **Light**<br>**Medium/Dark** | 195<br>284 | 25<br>32 | 1.14 (0.65-1.98)<br>**Ref** | 137<br>207 | 81<br>103 | 1.19 (0.83-1.71)<br>**Ref** | 164<br>245 | 44<br>67 | 0.98 (0.64-1.51)<br>**Ref** |
| **Skin response to sun exposure** | **Burnt/Blistered**<br>**Mild burn**<br>**Tan** | 130<br>157<br>184 | 16<br>17<br>22 | 1.03 (0.52-2.04)<br>0.91 (0.46-1.77)<br>**Ref** | 97<br>108<br>133 | 47<br>60<br>73 | 0.88 (0.56-1.39)<br>1.01 (0.66-1.55)<br>**Ref** | 106<br>142<br>154 | 34<br>30<br>44 | 1.12 (0.67-1.87)<br>0.74 (0.44-1.24)<br>**Ref** |

[a]Odds ratio, OR calculated for minor allele of each SNP in controls.
[b]Arsenic exposure adjusted for country.

**Supplementary table S8**. Telomere length associated SNPs and risk of BCC (dataset 2)

| **SNPs associated with Telomere Length** | | **Cases (N=522)** | **Controls (N=527)** | **OR (95% CI)[a]** | ***P*-value[b]** |
|---|---|---|---|---|---|
| **rs6060627 (*BCL2L1*)** | Major allele (C-allele)<br>Minor allele (T-allele)<br>missing | 376<br>144<br>2 | 386<br>134<br>7 | 1.00 (0.75-1.34) | 0.99 |
| **rs6772228 (*PXK*)** | Major allele (T-allele)<br>Minor allele (A-allele)<br>missing | 502<br>16<br>4 | 511<br>16<br>0 | 0.90 (0.43-1.86) | 0.77 |
| **rs9257445 (*ZNF311*)** | Major allele (G-allele)<br>Minor allele (C-allele)<br>missing | 393<br>111<br>18 | 377<br>145<br>5 | **0.73 (0.54-0.99)** | **0.04** |
| **rs1317082 (*TERC*)** | Major allele (A-allele)<br>Minor allele (G-allele)<br>missing | 389<br>125<br>8 | 393<br>129<br>5 | 1.03 (0.77-1.39) | 0.84 |
| **rs2487999 (*OBFC1*)** | Major allele (C-allele)<br>Minor allele (T-allele)<br>missing | 423<br>65<br>34 | 465<br>57<br>5 | 1.28 (0.86-1.90) | 0.22 |
| **rs7726159 (*TERT*)** | Major allele (C-allele)<br>Minor allele (A-allele)<br>missing | 359<br>161<br>2 | 339<br>181<br>7 | 0.85 (0.65-1.11) | 0.23 |
| **rs755017 (*RTEL1*)** | Major allele (A-allele)<br>Minor allele (G-allele)<br>missing | 456<br>50<br>16 | 477<br>49<br>1 | 1.03 (0.66-1.58) | 0.91 |
| **rs412658 (*ZNF676*)** | Major allele (C-allele)<br>Minor allele (T-allele)<br>missing | 325<br>189<br>8 | 309<br>201<br>17 | 0.84 (0.64-1.09) | 0.19 |
| **rs3027234 (*CTC1*)** | Major allele (C-allele)<br>Minor allele (T-allele)<br>missing | 440<br>82<br>0 | 406<br>104<br>17 | 0.71 (0.51-1.00) | 0.05 |
| **rs861539 (*XRCC3*)** | Major allele (C-allele)<br>Minor allele (T-allele) | 387<br>135 | 354<br>173 | **0.69 (0.52-0.91)** | **0.009** |

[a]Odds ratio, OR calculated in dataset 2 was adjusted for age, country and arsenic exposure.
[b]*P* -values were derived from $\chi^2$-test, two sided and considered statistically significant if < 0.05, indicated in bold.

**Supplementary Table S9.** β-estimates for SNP-telomere length association in controls and combined population (dataset 1) for one-sample study

| | | | Control population | | | | | | | Combined Population | | | | Genome-wide association studies | |
|---|---|---|---|---|---|---|---|---|---|---|---|---|---|---|---|
| **SNP** | **Chr. Position (hg19)** | **Gene** | **MAF** | **Allele (Major/minor)** | **N** | **F** | **β-estimate (standardized)[a]** | **SE** | ***P*** | **F** | **β-estimate (standardized)[a]** | **SE** | ***P*** | **ΔTL[b]** | **ref** |
| rs6060627 | 20:30262159 | *BCL2L1* | 0.28 | C/T | 520 | 11.37 | 0.236 | 0.003 | 0.29 | 24.74 | 0.338 | 0.002 | 0.05 | 36 | [1] |
| rs6772228 | 3:58376019 | *PXK* | 0.04 | T/A | 520 | 11.27 | -0.182 | 0.006 | 0.38 | 24.03 | ***0.010*** | 0.005 | 0.95 | -120 | [1] |
| rs9257445 | 6:28949206 | *ZNF311* | 0.29 | G/C | 518 | 10.55 | ***0.261*** | 0.002 | 0.23 | 22.76 | ***0.225*** | 0.002 | 0.19 | -38 | [1] |
| rs1317082 | 3:169497585 | *TERC* | 0.26 | A/G | 514 | 11.65 | -0.286 | 0.003 | 0.18 | 25.16 | -0.379 | 0.002 | 0.02 | -77 | [1] |
| rs2487999 | 10:105659826 | *OBFC1* | 0.13 | C/T | 520 | 11.40 | 0.014 | 0.003 | 0.95 | 24.99 | 0.082 | 0.002 | 0.61 | 100 | [1] |
| rs7726159 | 5:1282319 | *TERT* | 0.35 | C/A | 520 | 11.44 | 0.246 | 0.002 | 0.24 | 24.66 | 0.036 | 0.001 | 0.83 | 73 | [1] |
| rs755017 | 20:62421622 | *RTEL1* | 0.10 | A/G | 520 | 10.87 | ***0.113*** | 0.004 | 0.66 | 24.22 | ***0.157*** | 0.003 | 0.36 | -74 | [2] |
| rs412658 | 19:22359440 | *ZNF676* | 0.36 | C/T | 514 | 11.05 | ***0.209*** | 0.003 | 0.39 | 24.36 | ***0.176*** | 0.002 | 0.33 | -57 | [3] |
| rs3027234 | 17:8136092 | *CTC1* | 0.20 | C/T | 504 | 11.43 | -0.038 | 0.003 | 0.86 | 24.27 | ***0.162*** | 0.002 | 0.33 | -49 | [3] |
| rs861539 | 14:104165753 | *XRCC3* | 0.42 | G/A | 522 | 10.95 | 0.098 | 0.002 | 0.68 | 24.09 | -0.058 | 0.002 | 0.73 | n/a | |

[a]Standardized coefficient values from linear regression model after adjustment for age, country and log-transformed lifetime exposure of arsenic and 6 degrees of freedom (adj R square=0.11). The values correspond to equivalent age-related attrition of telomere for the corresponding SNP. The estimates not in accordance with published literature were highlighted in bold and italics. F, F-statistic values and SE, Standard error.

[b]Estimates of the per allele effect on leukocyte telomere length in base pairs as reported in genome-wide association studies.

| **Supplementary Table S10.** Mendelian randomization estimates for telomere length associated SNPs with BCC risk in one-sample study | | | | | |
|---|---|---|---|---|---|
| Using 10 SNPs (Nine telomere length associated SNPs and *XRCC3* polymorphism, rs861539) | | | | | |
| **Method** | **Estimate (95% CI)[a]** | **SE** | **OR (95% CI)[b]** | ***P*-value** | **Heterogeneity test statistic (*P*-value)** |
| ***Inverse-Variance Weighted*** | 0.60 (-0.02 to 1.23) | 0.32 | 1.83 (0.98-3.40) | 0.06 | 0.10 |
| ***Maximum-Likelihood*** | 0.61 (-0.02 to 1.23) | 0.32 | 1.83 (0.98-3.41) | 0.06 | |
| Using nine telomere length associated SNPs | | | | | |
| **Method** | **Estimate (95% CI)[a]** | **SE** | **OR (95% CI)[b]** | ***P*-value** | **Heterogeneity test statistic (*P*-value)** |
| ***Inverse-Variance Weighted*** | 0.50 (-0.04 to 1.04) | 0.28 | 1.65 (0.96-2.83) | 0.07 | 0.30 |
| ***Maximum-Likelihood*** | 0.50 (-0.04 to 1.04) | 0.28 | 1.65 (0.96-2.83) | 0.07 | |

[a]Mendelian randomization estimates for short telomeres on BCC risk.

[b]Odds Ratio, OR and 95% confidence interval, CI were calculated by taking the exponential of the estimates. SE, standard error.
